# ShinyGPAS: Interactive genomic prediction accuracy simulator based on deterministic formulas

**DOI:** 10.1101/164772

**Authors:** Gota Morota

## Abstract

**Background:** Deterministic formulas highlight the relationships among prediction accuracy and potential factors influencing prediction accuracy prior to performing computationally intensive cross-validation. Visualizing such deterministic formulas in an interactive manner may lead to a better understanding of how genetic factors control prediction accuracy.

**Results:** The software to simulate deterministic formulas for genomic prediction accuracy was implemented in R and encapsulated as a web-based Shiny application. ShinyGPAS (Shiny Genomic Prediction Accuracy Simulator) simulates various deterministic formulas and delivers dynamic scatter plots of prediction accuracy vs. genetic factors impacting prediction accuracy, while requiring only mouse navigation in a web browser. ShinyGPAS is available at: <u>https://chikudaisei.shinyapps.io/shinygpas/</u>.

**Conclusion:** ShinyGPAS is a shiny-based interactive genomic prediction accuracy simulator using deterministic formulas. It can be used for interactively exploring potential factors influencing prediction accuracy in genome-enabled prediction, simulating achievable prediction accuracy prior to genotyping individuals, or supporting in-class teaching. ShinyGPAS is open source software and it is hosted online as a freely available web-based resource with an intuitive graphical user interface.

## Background

Prediction of genomic values from high-dimensional single nucleotide polymorphisms is a primal interest in animal breeding and quantitative genetics (Meuwissen et al., 2001; Goddard, 2017). A deterministic formula such as the one proposed by Daetwyler et al. (2008) highlights the relationship between prediction accuracy and potential factors influencing prediction accuracy. In general, deterministic formulas compute expected predictive correlation (or prediction *R*^2^ of phenotypes) on the basis of a number of factors that are potentially useful to assess prediction accuracy before performing computationally demanding crossvalidation (CV). It also allows us to decide the optimal design for training populations. Not only theoretical derivations of deterministic formulas but also their applications are active research areas. For instance, Brard and Ricard (2015) recently performed comparison and meta-analysis of deterministic formulas. Erbe et al. (2013) inferred parameters that influence prediction accuracy in deterministic formulas via a maximum likelihood. Collectively, these studies have shed new light on alternative aspects of factors influencing predictive performance that may not be obvious from empirical genome-enabled prediction analysis based on CV.

In particular, visualizing such deterministic formulas may lead to a better understanding of how genetic factors control prediction accuracy. Typically, visualization involves generating a static two-dimensional graph, where the y-axis is the genomic prediction accuracy and the x-axis is one of the factors influencing prediction accuracy, while keeping the other factors constant. Given that this type of static graph is a snapshot of complex dynamic system, if users want to change parameters, they need to re-type and re-execute the code. To overview the whole landscape of genomic prediction simulation, we need an efficient visualization tool that is capable for generating interactive as well as dynamic graphs. The objective of this article is to describe a Shiny-based web application called ShinyGPAS (Shiny Genomic Prediction Accuracy Simulator), which produces interactive graphs and offers an intuitive graphical user interface (GUI) for simulating genomic prediction accuracy based on deterministic formulas.

## Software description

### Overview of software architecture

ShinyGPAS is implemented entirely in R, which is an open source programming language and environment for performing statistical computing and data visualization (R Core Team, 2017). The GUI is provided by the shiny R package (Chang et al., 2017), a web application framework for R. ShinyGPAS is a Shiny application that leverages R and the shiny package to construct an intuitive framework for deterministic formulas using dynamic interaction and visualization. The ShinyGPAS user interface is shown in Figure 1. Although ShinyGPAS is R-based software, it does not require users to either be familiar with the programming language nor download the software on a local computer. The underlying R code is encapsulated by Shiny and offered as cohesive web-based software to be usable solely by mouse navigation in a web browser. This increases accessibility to the software, especially for users with less R programming experience. ShinyGPAS is deployed through the cloud-based shinyapps.io platform for hosting Shiny web applications (<u>https://www.shinyapps.io/</u>).

**Figure 1:**
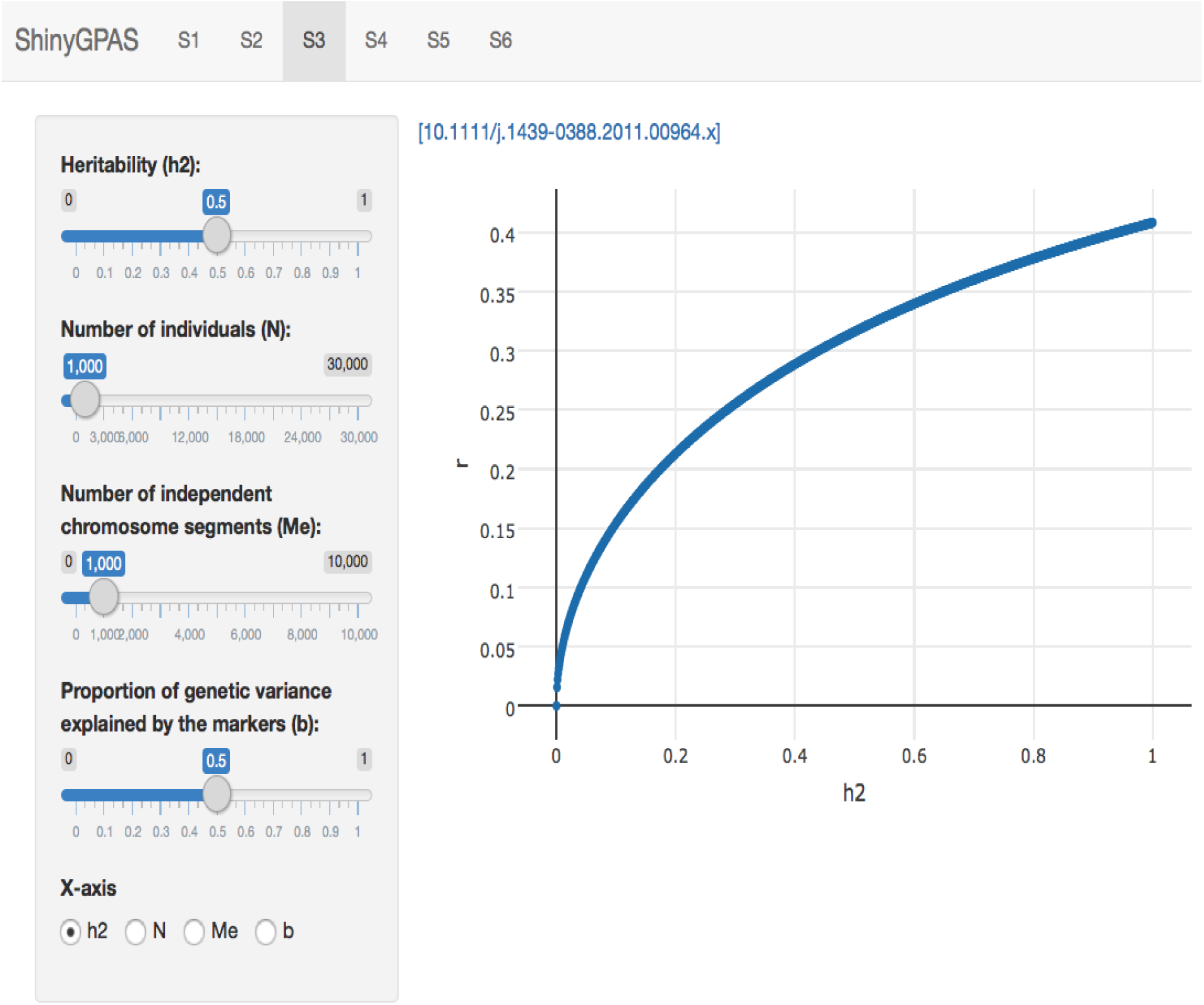
Each deterministic formula is implemented in a tab on the top. The y-axis is the prediction accuracy or squared prediction accuracy and the x-axis is one of the parameters. Parameters including heritability, the number of individuals, the number of independent chromosome segments, effective population size, and the proportion of genetic variance explained by the markers can be set by the user.

### Deterministic formulas

ShinyGPAS currently delivers six simulators based on deterministic formulas described in a) Daetwyler et al. (2008, 2010), b) Goddard (2009), c) Goddard et al. (2011), d) de los Campos et al. (2013), e) Karaman et al. (2016), and f) Wientjes et al. (2016). The first five formulas predict accuracy or squared prediction accuracy within populations whereas the last one is designed for multipopulation including multienvironment and multitrait scenarios. Deterministic formulas are functions derived from the combinations of the number of individuals in a reference set, the number of independent chromosome segments underlying the trait, the effective population size, the proportion of genetic variance explained by the molecular markers, and heritability. Shiny-based interactive application offers the implementation of dynamic deterministic formulas, allowing to evaluate the simultaneous impact of all the parameters described above on the degree of prediction accuracy. A user can click a link located within each deterministic formula simulator to access original journal articles. Below are deterministic formulas currently implemented in ShinyGPAS.

- Daetwyler et al. (2008, 2010)

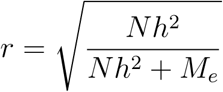

where *N* is the number of individuals, *h*^2^ is the heritability, and *M*_*e*_ is the number of independent chromosome segments.
- Goddard (2009)

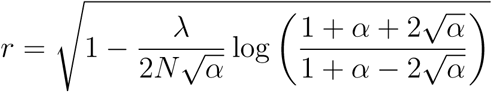

where *λ* is *M*_*e*_*/*(*h*^2^ log(2*N*_*e*_)), *α* is 1 + 2(*M*_*e*_*/Nh*^2^ log(2*N*_*e*_)), and *N*_*e*_ is the effective population size.
- Goddard et al. (2011)

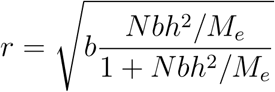

where *b* is the proportion of genetic variance explained by the markers.
- de los Campos et al. (2013)

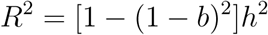

where *b* is the average regression coefficient of the marker-based genomic relationships on causal loci derived genomic relationships.
- Karaman et al. (2016)

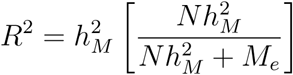

where 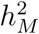 is the genomic heritability, which is the proportion of phenotypic variance that is explained by regression on markers.
- Wientjes et al. (2016)

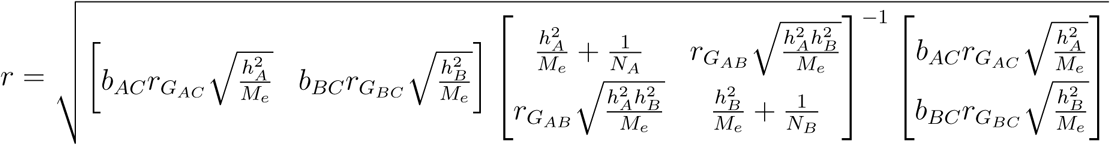

where *b*_*AC*_ is the square root of the proportion of the genetic variance in predicted population C explained by the markers in training population A, *r*_*G_AC_*_ is the genetic correlation between populations A and C, 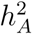is heritability in population A, *b*_*BC*_ is the square root of the proportion of the genetic variance in predicted population C explained by the markers in training population B, *r*_*G_BC_*_ is the genetic correlation between populations B and C, 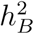 is heritability in population B, *N*_*A*_ is the number of individuals in population A, *N*_*B*_ is the number of individuals in population B, and *r*_*GAB*_ is the genetic correlation between populations A and B. This deterministic formula combines two populations A and B to predict prediction accuracy in population C (Wientjes et al., 2016).

### Program input

A typical workflow starts from selecting one of the tab panels on the top (Figure 1) and then moving to a preferred deterministic formula simulator. Each of deterministic formula captures a different aspect of the genotype-phenotype map in the context of genomic prediction accuracy. Thus, navigating interactively visualized deterministic formulas may highlight the common patterns as well as differences among them. A suite of available parameters such as *h*^2^, *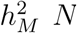, M*_*e*_, *N*_*e*_, and *b* are located in the sidebar panel. Shiny slider provides possible input values one can choose from pre-defined ranges. Users can pick a preferred value by a simple mouse navigation. A radio button located on the bottom offers possible options for factors influencing prediction accuracy to be used to determine the x-axis. The Shiny reactive expressions are utilized in ShinyGPAS to efficiently cache results and ease computational burden to ensure high speed of processing during an interactive session.

### Program output

Rendering interactive graphs from deterministic formulas are achieved by the plotly R package (Sievert et al., 2017). The main engine plotly.js, which is built on top of JavaScript and the visualization library D3.js, was used to create a scatter plot. The y-axis is pre-fixed with prediction accuracy (*r*) or squared prediction accuracy(*R*^2^). Users can choose the x-axis from one of the parameters including *h*^2^,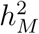, *N, M*_*e*_, *N*_*e*_, or *b*. A scatter plot is dynamically updated when users vary slider input values of factors influencing prediction accuracy. The plotly.js generates a scatter plot with a toolbar coupled with useful zooming in and zooming out capabilities. Also, hovering the mouse pointer over a specific point of plot shows the exact values of x and y axes. A multipopulation genomic prediction simulation is enabled by the plotly 3D scatter plot functionality, where x and y axes take parameters from two training populations and z-axis shows prediction accuracy. Rotating the 3D scatter plot is possible around all x, y and z axes to inspect prediction accuracy from different surfaces. In addition, the toolbar provides features such as download button, box select, lasso select, autoscale, reset, and toggle spike lines features for interactivity. ShinyGPAS is available at: <u>https://chikudaisei.shinyapps.io/shinygpas/</u>.

## Conclusions

A Shiny application has great potential to deliver interactive data analysis and visualization in a web browser. Yet there is limited application of this type of tool in animal breeding and quantitative genetics. The use of Shiny framework allows users to convert deterministic formulas of genomic prediction accuracy into interactive graphics in an engaging and straightforward manner. ShinyGPAS can be used for interactive exploration of potential factors influencing prediction accuracy in genome-enabled prediction, simulation of achievable prediction accuracy prior to genotyping individuals, or supporting in-class teaching. The ShinyGPAS source code has been made publicly available on GitHub: <u>https://github.com/morota/ShinyGPAS</u>.

## Declarations

### Authors’ contributions

GM designed and developed the software, and wrote the manuscript.

## References

Brard, S. and Ricard, A. (2015). Is the use of formulae a reliable way to predict the accuracy of genomic selection? Journal of Animal Breeding and Genetics, 132(3):207–217.

Chang, W., Cheng, J., Allaire, J., Xie, Y., and McPherson, J. (2017). shiny: Web Application Framework for R. R package version 1.0.3.

Daetwyler, H. D., Pong-Wong, R., Villanueva, B., and Woolliams, J. A. (2010). The impact of genetic architecture on genome-wide evaluation methods. Genetics, 185(3):1021–1031.

Daetwyler, H. D., Villanueva, B., and Woolliams, J. A. (2008). Accuracy of predicting the genetic risk of disease using a genome-wide approach. PloS one, 3(10):e3395.

de los Campos, G., Vazquez, A. I., Fernando, R., Klimentidis, Y. C., and Sorensen, D. (2013). Prediction of complex human traits using the genomic best linear unbiased predictor. PLoS Genet, 9(7):e1003608.

Erbe, M., Gredler, B., Seefried, F. R., Bapst, B., and Simianer, H. (2013). A function accounting for training set size and marker density to model the average accuracy of genomic prediction. PLoS One, 8(12):e81046.

Goddard, M. (2009). Genomic selection: prediction of accuracy and maximisation of long term response. Genetica, 136(2):245–257.

Goddard, M. (2017). Can we make genomic selection 100% accurate? Journal of Animal Breeding and Genetics, 134(4):287–288.

Goddard, M., Hayes, B., and Meuwissen, T. (2011). Using the genomic relationship matrix to predict the accuracy of genomic selection. Journal of Animal Breeding and Genetics, 128(6):409–421.

Karaman, E., Cheng, H., Firat, M. Z., Garrick, D. J., and Fernando, R. L. (2016). An upper bound for accuracy of prediction using GBLUP. PloS one, 11(8):e0161054.

Meuwissen, T. H., Hayes, B. J., and Goddard, M. E. (2001). Prediction of total genetic value using genome-wide dense marker maps. Genetics, 157(4):1819–1829.

R Core Team (2017). R: A Language and Environment for Statistical Computing. R Foundation for Statistical Computing, Vienna, Austria.

Sievert, C., Parmer, C., Hocking, T., Chamberlain, S., Ram, K., Corvellec, M., and Despouy, P. (2017). plotly: Create Interactive Web Graphics via ‘plotly.js’. R package version 4.6.0.

Wientjes, Y. C., Bijma, P., Veerkamp, R. F., and Calus, M. P. (2016). An equation to predict the accuracy of genomic values by combining data from multiple traits, populations, or environments. Genetics, 202(2):799–823.

